# Structural Insights into Pink-eyed Dilution Protein (Oca2)

**DOI:** 10.1101/2022.12.09.519718

**Authors:** Shahram Mesdaghi, David L. Murphy, AJ Simpkin, Daniel J. Rigden

## Abstract

Recent innovations in computational structural biology have opened an opportunity to revise our current understanding of the structure and function of clinically important proteins. This study centres on human Oca2 which is located on mature melanosomal membranes. Mutations of Oca2 can result in a form of oculocutanous albinism which is the most prevalent and visually identifiable form of albinism. Sequence analysis predicts Oca2 to be a member of the SLC13 transporter family but it has not been classified into any existing SLC families. The modelling of Oca2 with AlphaFold2 and other advanced methods shows that, like SLC13 members, it consists of a scaffold and transport domain and displays a pseudo inverted repeat topology that includes re-entrant loops. This finding contradicts the prevailing consensus view of its topology. In addition to the scaffold and transport domains the presence of a cryptic GOLD domain is revealed that is likely responsible for its trafficking from the endoplasmic reticulum to the Golgi prior to localisation at the melanosomes and possesses known glycosylation sites. Analysis of the putative ligand binding site of the model shows the presence of highly conserved key asparagine residues that suggest Oca2 may be a Na^+^/dicarboxylate symporter. Known critical pathogenic mutations map to structural features present in the repeat regions that form the transport domain. Exploiting the AlphaFold2 multimeric modelling protocol in combination with conventional homology modelling allowed the building of a plausible homodimer in both an inward- and outward-facing conformation supporting an elevator-type transport mechanism.

## Introduction

Albinism is a hereditary condition affecting the synthesis of melanin. The most prevalent and visually identifiable form of albinism is oculocutanous albinism. Oculocutanous albinism is a recessive disorder where individuals have a phenotype exhibiting melanin deficiency in the skin, hair, and eyes. Oculocutanous albinism results from mutations in genes that code for proteins that are involved in melanin production. The gene affected is used to classify the type of oculocutanous albinism into one of the 7 subtypes (oculocutanous albinism 1-7); oculocutanous albinism type 1:TYR, oculocutanous albinism type 2:OCA2, oculocutanous albinism type 3:TYRP1, oculocutanous albinism type 4:SLC45A2, oculocutanous albinism type 6:SLC24A5, oculocutanous albinism type 7:LRMDA and oculocutanous albinism type 5 gene is located on chromosome 4q24 (Yang et al., 2019). Accurate diagnosis of the sub-type can only be achieved by a genetic screen (Arveiler, Lasseaux, & Morice-Picard, 2017). The most prevalent form of oculocutanous albinism is oculocutanous albinism type 2 in which mutations in the OCA2 gene cause changes in the transmembrane protein p protein (Oca2) thereby impacting melanin production. Polymorphisms of the OCA2 gene have been shown to be major contributor to skin colour (Lao, de Gruijter, van Duijn, Navarro, & Kayser, 2007) and are thought to underlie blue eye colour in humans (Eiberg et al., 2008). Oca2 is expressed in melanocytes and retinal pigment epithelium (RPE) where it is restricted to melanosomes.

Melanosomes are “lysosome-related organelles” but are functionally and morphologically distinct from lysosomes as they have an acidic luminal pH (Griffiths, 2002) and possess cell– type-specific cargo proteins (Raposo, Marks, & Cutler, 2007). Trafficking pathways deliver these cargo proteins to immature melanosomes which contributes to their maturation (Raposo & Marks, 2007). Oca2 is located in the mature melanosomal membrane where it has been shown to control chloride conductance across the lipid bilayer (Bellono, Escobar, Lefkovith, Marks, & Oancea, 2014). This chloride conductance is coupled to proton motive force and is related to maintenance of the optimal luminal pH for the tyrosinase function involved in production of melanin (Bellono et al., 2014) be. The currently accepted model of Oca2, based on hydrophobicity profiles, describes Oca2 as a 12 transmembrane helix protein with two luminal loops and an N-terminal disordered cytoplasmic loop (Gardner et al., 1992). Oca2 is glycosylated in the N-terminal luminal loop and the N-terminal cytoplasmic loop of Oca2 possesses dileucine motifs; both of these features are important for the trafficking of Oca2 from the ER to the melanosomes through a series of intracellular compartments (Sitaram et al., 2009).

This study employs deep learning modelling methods to argue for a revised topology for Oca2. Deep learning methods such as DMPfold (Greener, Kandathil, & Jones, 2019), trRosetta (Bassot & Elofsson, 2021) and AlphaFold2 (Jumper et al., 2021) build predicted protein structures by predicting inter residue distances, main chain hydrogen bond network and torsion angles and utilizing these as restraints in the model building process. Benchmarking these methods have demonstrated that they work just as well for membrane proteins as they do for soluble proteins (Greener et al., 2019; Hegedűs, Geisler, Lukács, & Farkas, 2022). DMPfold was shown to be able to model transmembrane proteins with a TM-score of at least 0.5 to the native structure and obtain a mean TM-score of 0.74 (Greener et al., 2019). The accuracy of AlphaFold2 transmembrane protein modelling has been tested by exploring the construction of structures from the ABC protein superfamily. For these transmembrane proteins AlphaFold2 performed exceedingly well when testing template-free structure prediction as well as attempting a new ABC fold, dimer modelling, and stability in molecular dynamics simulations (Hegedűs et al., 2022).

The modelling of Oca2 using AlphaFold2 predicts the presence of a pseudo inverted repeat that forms a pore region flanked with two highly conserved re-entrant loops. Additionally, a luminal loop proceeding the first transmembrane helix is predicted to be GOLD-like domain that allows trafficking through the Golgi from the endoplasmic reticulum to finally localise at the melanosomal membrane. The newly proposed topology shares features with sodium-carboxylate transporters (NaCT) which is supported by the obvious sequence homology.

## Methods

### Hardware

Local ColabFold model building was performed on an Ubuntu 18.04.6 workstation AMD Ryzen Threadripper 2990WX 32 Core CPU (3.0GHz) with 64GB RAM. GPU acceleration was performed by an ASUS TUF GeForce RTX 3080 OC LHR 12GB GDDR6X Ray-Tracing Graphics Card, 8960 Core, 1815MHz Boost.

### Pfam database screening

Searches using the sequence of Oca2 were made against the Pfam-A_v35.0 (RRID:SCR_004726) (El-Gebali et al., 2019) database using the HHPred (RRID:SCR_010276) v3.0 server (Zimmermann et al., 2018) with default parameters (-p 20 -Z 10000 -loc -z 1 -b 1 -B 10000 -ssm 2 -sc 1 -seq 1 -dbstrlen 10000 -norealign -maxres 32000 -contxt /cluster/toolkit/production/bioprogs/tools/hh-suite-build-new/data/context_data.crf) and eight iterations for MSA generation in the HHblits (Zimmermann et al., 2018) stage.

### Structural database screening

Dali (RRID:SCR_013433) v5.0 server (Holm & Laakso, 2016) was used to screen the PDB (Burley et al., 2018) and the AF human proteome database (David, Islam, Tankhilevich, & Sternberg, 2022) for structural homologues of Oca2. Pairwise alignments were also performed by the Dali server.

### Model building

An initial Oca2 model was obtained from the AFDB (David et al., 2022). The construction of the inward-facing homodimeric form and attempted alternate conformations was performed by a local installation of ColabFold (Mirdita et al., 2022).

The outward-facing monomers were constructed by first building a homology model of Oca2 using an outward facing structure of a homologue. The HHpred server was used to identify 6wtw as a close homologue (99.97% probability). Modeller (Eswar, Eramian, Webb, Shen, & Sali, 2008) functionality of the MPI bioinformatics toolkit server was used to build the homology model. The homology model as a template along with custom MSAs of varying depths were used as inputs in a local installation of ColabFold. Five models at each MSA depth were constructed and the model with the highest mean pLDDT score was selected for examination.

The outward facing homodimer was constructed using a local installation of ColabFold with the Modeller homology structure used as a template and an MSA with depth of 15 sequences.

### MSAs

MSAs were build using the HHblits server (Zimmermann et al., 2018) using default settings. The reduction in MSA depth, as a strategy to assist exploration of conformational diversity in AF outputs, was achieved by randomly selecting sequences from the HHblits output.

### Docking

The Webina server (Kochnev, Hellemann, Cassidy, & Durrant, 2020) utilizing Autodock (Trott & Olson, 2009) was used to dock citrate into the putative binding pocket of Oca2. A docking box size of 35×35×35 was used with the default coordinates for the box center. Prodigy (Xue, Rodrigues, Kastritis, Bonvin, & Vangone, 2016) was used to perform docking rescoring.

## Results and Discussion

### Oca2 is a member of the IT Superfamily

Oca2 is an 838-residue transmembrane protein annotated in UniProt with the Pfam domain CitMHS (PF03600). An HHpred search of the Pfam database reveals that Oca2 also possesses strong sequence similarity to other members of the Ion Transporter (IT) Superfamily with HHpred probability scores above 99.9% (Table 1).

**Table 1:** HHpred results for screen of Oca2 sequence against Pfam

| Pfam Accession | Pfam Name | Probability | E-Value | Query HMM | Template HMM |
| --- | --- | --- | --- | --- | --- |
| PF02040.18 | ArsB | 100 | 1.90E-26 | 334-831 | 2-422 |
| PF00939.22 | Na_sulph_symp | 100 | 3.50E-25 | 324-837 | 23-473 |
| PF06450.15 | NhaB | 100 | 1.20E-24 | 328-831 | 42-503 |
| PF16980.8 | CitMHS_1 | 99.9 | 2.30E-23 | 332-829 | 11-442 |
| PF07399.14 | Na_H_antiport_2 | 99.9 | 1.50E-22 | 328-826 | 1-413 |
| PF06808.15 | DctM | 99.9 | 6.00E-20 | 339-830 | 1-410 |
| PF02447.19 | GntP_permease | 99.9 | 9.70E-19 | 329-834 | 1-443 |
| PF03600.19 | CitMHS | 99.9 | 8.00E-19 | 342-773 | 1-348 |

ArsB protein is established as a bacterial arsenite efflux pump (Rosen, 1999). Although it is common for ArsB to complex with the ATPase ArsA to form an ATP-driven pump that expels arsenite (Dey & Rosen, 1995) it has been shown that ArsB can function independently as an arsenite efflux pump by coupling with proton motive force (Ji, Bacteriology, & 1992, n.d.). Although Oca2 shows no sequence similarity with Cl^-^ transporters it has been demonstrated experimentally that Oca2 is required for melanosomal anion efflux contributing to the mediation of chloride-selective anion conductance which in turn modulates melanosome pH thereby regulating melanin synthesis (Bellono et al., 2014).

### SLC13 members have a pseudo inverse repeat topology

The IT Superfamily is made up of both symporters and antiporters (Sauer et al., n.d.). Experimental structures are available for some members, but no experimental structures are available for Oca2. To identify experimental structures of close evolutionary relatives to Oca2, the sequence was screened against the PDB using HHpred. There were three hits above 99.9% probability comprising members of Pfam families DASS (divalent anion sodium symporter) family sodium-coupled anion symporter, Solute carrier family 13 member 5 and NaDC (Table 2). Additionally, the AlphaFold2 model of Oca2 was screened against the full PDB using Dali. All hits above a Z-score of 35 were Na^+^ symporters in the inward conformation with the top hit being 7jsj, the Solute carrier family 13 member 5 and NadC (Supplementary Table 1). Furthermore, the AlphaFold2 model of Oca2 was screened against the Alphafold database of the human proteome using Dali; there were five hits above a Z-score of 30 for the conserved Pfam domain (PF16980); Solute Carrier Family 13 Members 1,2,3,4 and 5 (Table 3).

**Table 2:** HHpred results for screen of Oca2 sequence against PDB

| DBAccession | Pfam Name | Probability | E-Value | Query HMM | Template HMM |
| --- | --- | --- | --- | --- | --- |
| 6wtw_B | DASS family sodium-coup | 100 | 5.00E-24 | 323-837 | 32-490 |
| 7jsk_B | Solute carrier fam 13 mem 5 | 99.99 | 8.20E-24 | 327-836 | 31-552 |
| 5ul9_D | Transporter, NadC family | 99.9 | 3.70E-23 | 326-832 | 23-444 |

**Table 3:** Dali results for structural screen (residues 331-831) of Oca2 against Human AlphaFold database. Light Tan: Hits against Oca2 conserved Pfam (PF16980) domain; Dark tan: hits against Oca2 N-terminal luminal loop

| Name | Z-Score |
| --- | --- |
| Human:Af-Q86yt5-F1 Solute Carrier Family 13 Member 5; | 36 |
| Human:Af-Q9ukg4-F1 Solute Carrier Family 13 Member 4; | 34.6 |
| Human:Af-Q8wwt9-F1 Solute Carrier Family 13 Member 3; | 34.1 |
| Human:Af-Q13183-F1 Solute Carrier Family 13 Member 2; | 33.4 |
| Human:Af-Q9bzw2-F1 Solute Carrier Family 13 Member 1; | 29.7 |
| Human:Af-O00337-F1 Sodium/Nucleoside Cotransporter 1; | 10.6 |
| Human:Af-Q9has3-F1 Solute Carrier Family 28 Member 3; | 10.2 |
| Human:Af-O43868-F1 Sodium/Nucleoside Cotransporter 2; | 10.3 |
| Human:Af-A6nh21-F1 Serine Incorporator 4; | 9.1 |

Solute carrier (SLC) proteins are integral membrane transport proteins that are classified into 66 families (Hediger et al., 2004; Perland & Fredriksson, 2017). Members within each family have greater than 20% sequence identity. However, the homology between solute carrier families maybe non-existent (Hoglund, Nordstrom, Schioth, & Fredriksson, 2011) as the basis for the introduction of a family as a solute carrier protein is related to functionality rather than an evolutionary link. Currently there is one structure available for a mammalian SLC13 protein; 562 residue long sodium-dependent citrate transporter (NaCT), SLC13 member 5. NaCT displays inverted repeat pseudo-symmetry relating the N-terminal half to the C-terminal half (Sauer et al., 2021) with each repeat containing a re-entrant loop packing against a broken transmembrane helix, followed by a cytosolic amphipathic helix parallel to the membrane plane, a second re-entrant loop packing against a broken transmembrane helix then finally a transmembrane helix (Figure 1).

**Figure 1:**
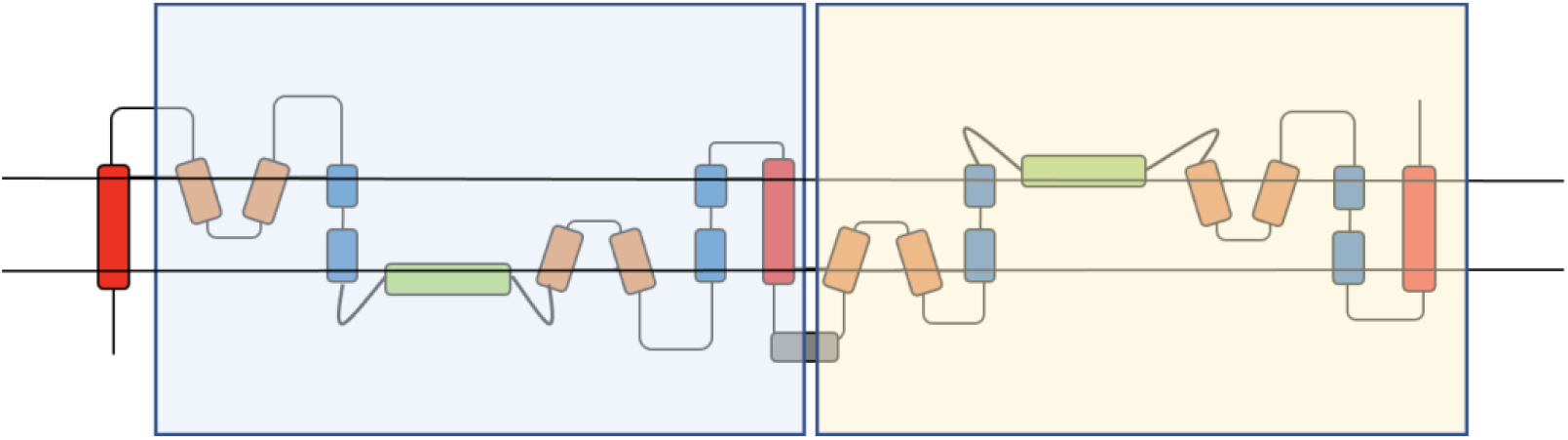
NaCT topology. Shaded regions highlight the pseudo inverse repeat. Red: transmembrane helix; Orange: re-entrant loop; Blue: broken helix; Green: amphipathic helix; Grey: extra-membrane helix.

### Oca2 has a pseudo inverse repeat topology

Examination of the Oca2 AlphaFold2 model (Figure 2) reveals a more complex topology compared to the currently accepted model of 12 transmembrane helices with two luminal loops and an N-terminal disordered cytoplasmic loop (Figure 4).

**Figure 2:**
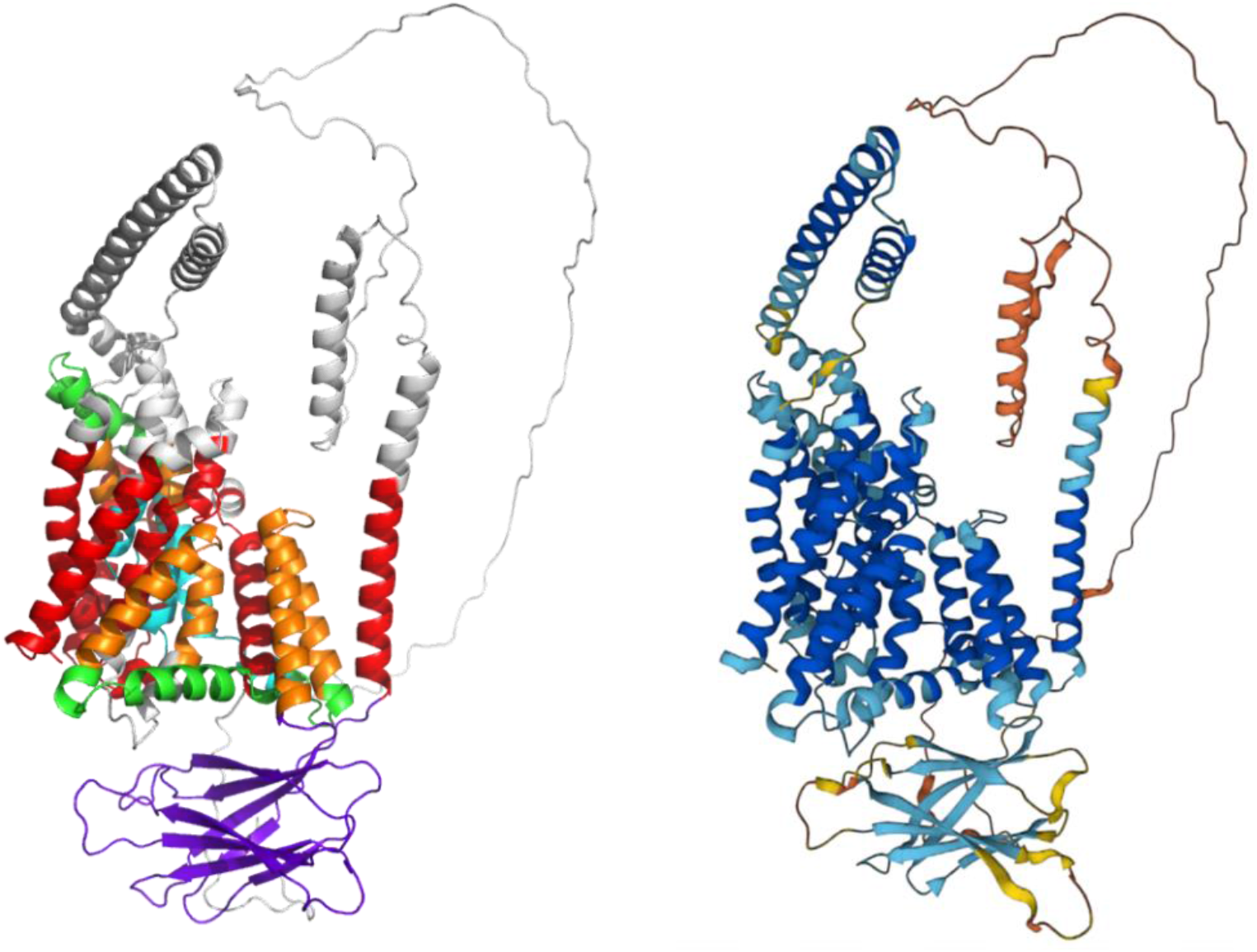
AlphaFold2 ca2 model. a) Red: transmembrane helix; Orange: re-entrant loop; Blue: broken helix; Green: amphipathic helix; Purple: ‘N-terminal luminal loop’-putative GOLD domain; Grey: extra-membrane helix. b) Coloured by AlphaFold2 per-residue confidence score (pLDDT) between 0 and 100. pLDDT>90 (blue) to pLDDT<50 (red).

**Figure 3:**
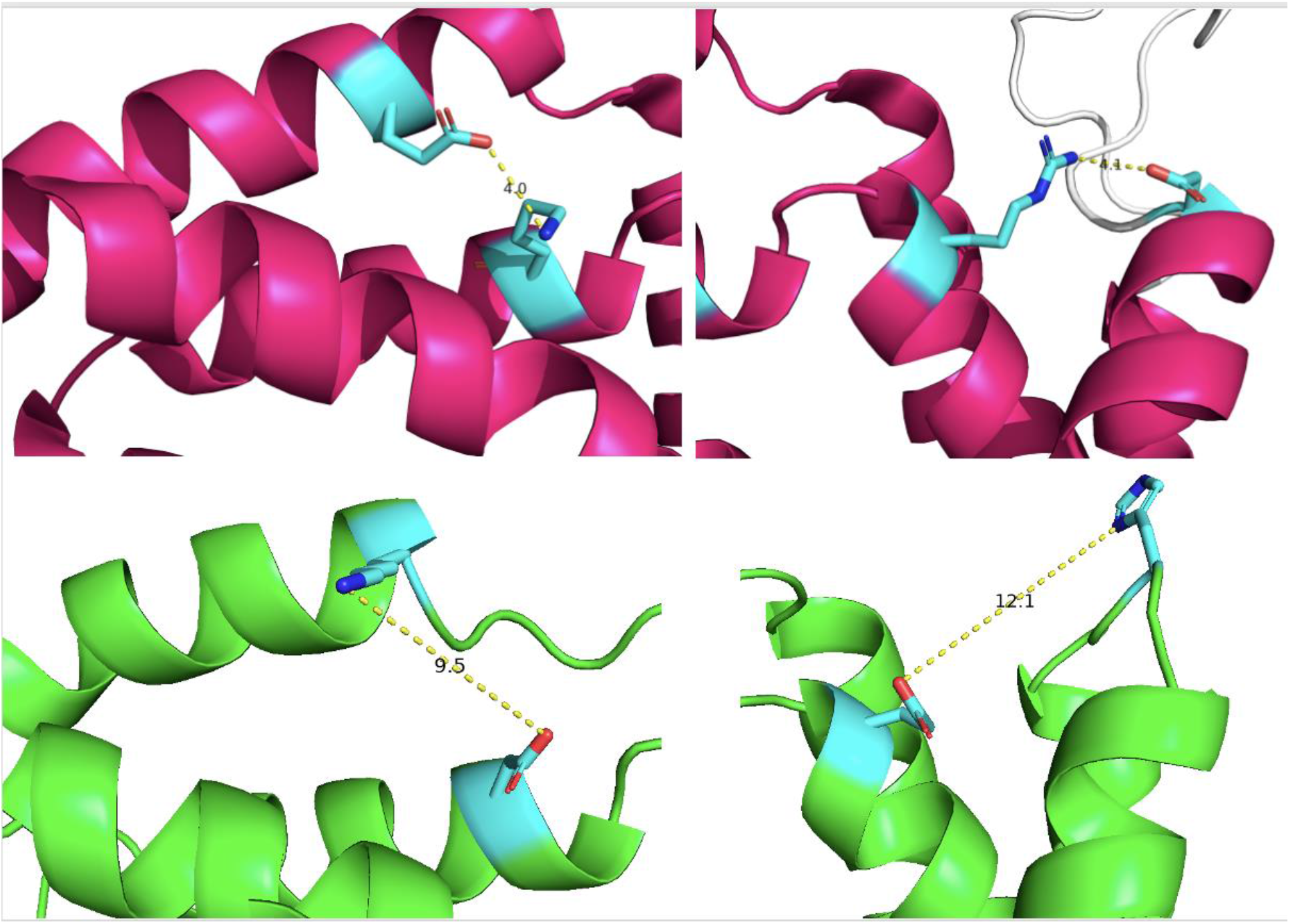
Presence of possible salt bridges stabilising the scaffold domain. Magenta: NaCT salt bridges; Left: Lys107 – Glu305, Right: Arg102 – Asp398. Green: Oca2 possible salt bridges; Left: Asp408 - Lys614, Right: Glu403 with His668.

**Figure 4 Top:**
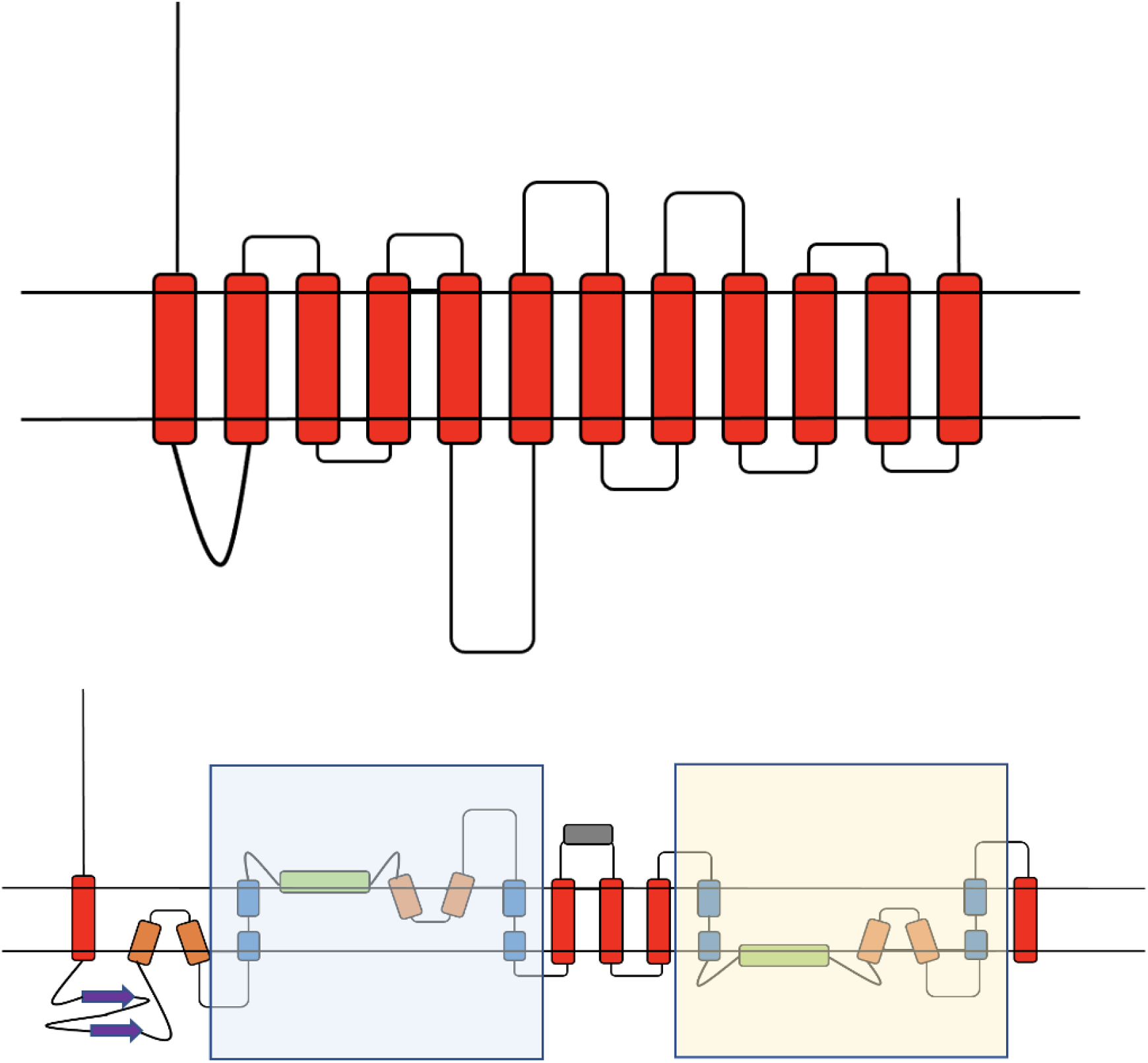
Current consensus view of Oca2 topology (Sitaram et al., 2009). Bottom: Proposed Oca2 Topology. Shaded regions are the pseudo inverse repeat. Red: transmembrane helix; Orange: re-entrant loop; Blue: broken helix; Green: amphipathic helix; Purple: ‘N-terminal luminal loop’; Grey: extra-membrane helix.

The AlphaFold2 model shares topological features with NaCT: a pseudo inverse repeat, each possessing a broken transmembrane helix, an amphipathic helix and a re-entrant loop packing against a broken transmembrane helix. Extrapolating functional annotations from homologues of Oca2 that have an experimental structure identified from the HHpred PDB screen indicates that each repeat unit of Oca2 possesses a transport domain made up of a re-entrant loop packed with transmembrane helix, where the transmembrane helix is broken in the centre. The amphipathic helices of each unit link the transport domain to a scaffold domain formed by the other helices of the conserved CitMHS domain. It has been shown in the experimental homologues that during the transport cycle, the two amphipathic helices are fixed in space with respect to the scaffold domain and cradle the transport domain during the conformational transition between the outward- and inward-facing states (Drew & Boudker, 2016; Sauer et al., n.d.). The rigidity of the Oca2 amphipathic helices could be achieved by salt bridges, between Glu403 with His668 as well as Asp408 with Lys614. Indeed, mutations of the equivalent residues (Arg102 – Asp398 and Lys107 – Glu305), in NaCT results in the abolition of substrate transport (Sauer et al., 2021). The distances between the potential salt bridge forming residues in the Oca2 model are beyond the 4Å threshold distance for salt bridge formation although in both cases the residues are located adjacent to flexible disordered loops which may facilitate the potential interaction for the residues forming a salt bridge (Figure 3).

Further inspection of the model identified additional structural features outside of the repeating units; a 171 residue long cytosolic N-terminal disordered region, a luminal 130 residue forming an eight stranded beta sandwich and a 95-residue cytosolic helical region separating the two inverse repeats (Figure 5).

**Figure 5:**
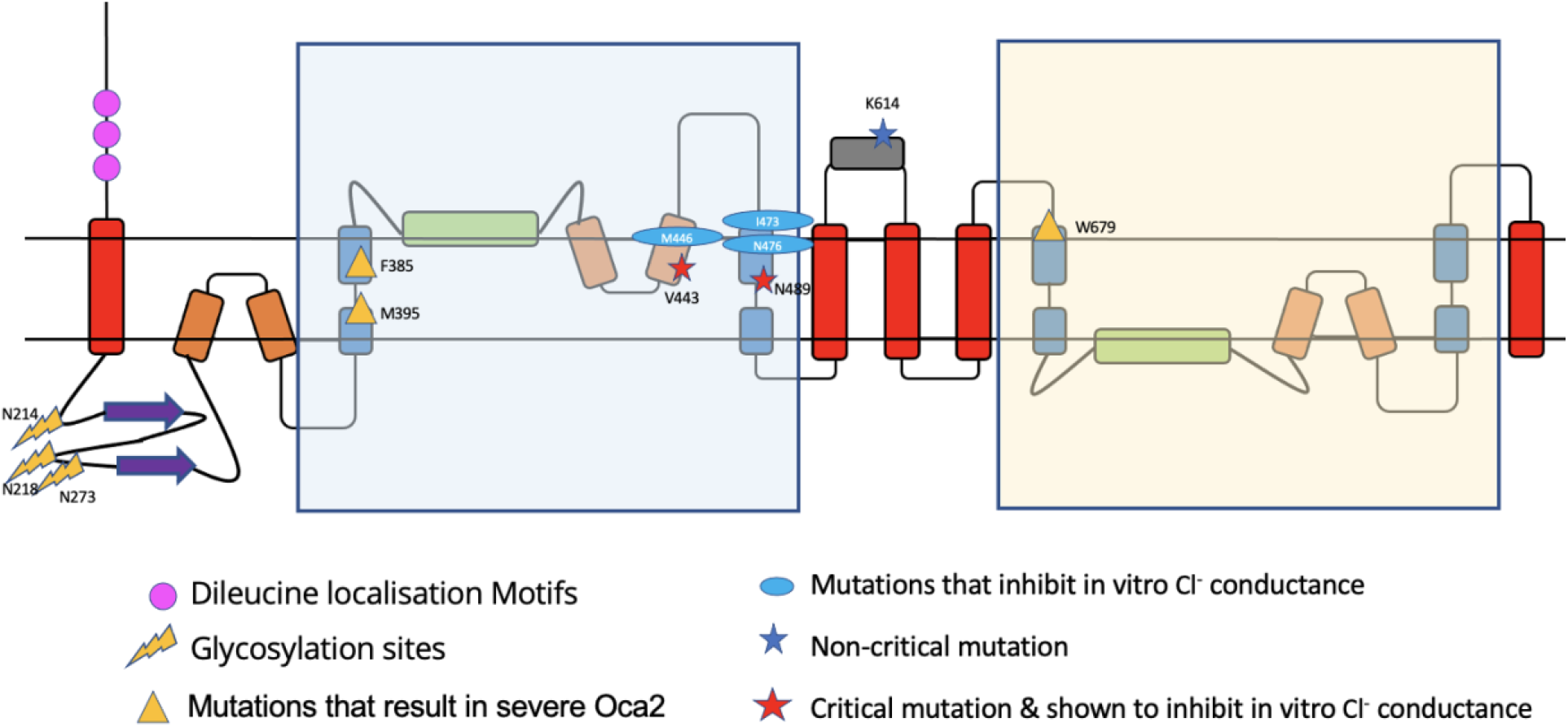
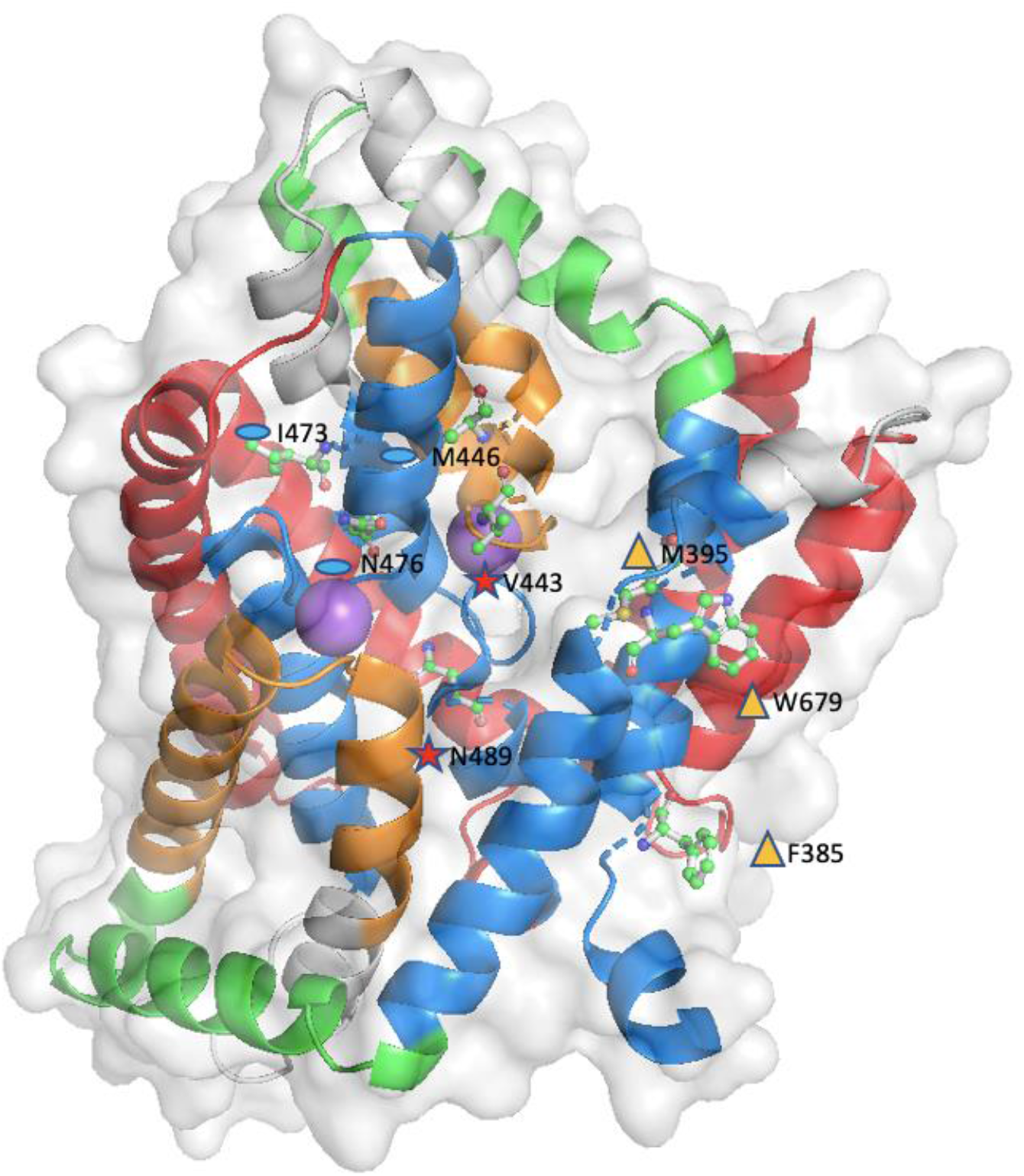
a) Oca2 topology with trafficking and example mutation sites mapped. b) Example mutations mapped on to the AlphaFold2 Oca2 structure (ball and stick); Na^+^ ions shown as purple spheres.

### Oca2 dileucine motifs responsible for melanosome localisation are located on the disordered cytosolic N-terminal region

The model of Oca2 has a N-terminal disordered loop. AlphaFold2 models this loop packed with transmembrane helices and when placed into a membrane bi-layer using the OMP server the 170 residue long cytosolic N-terminal disordered region crosses the membrane which is an obvious error. The region has very low confidence (pLDDT < 50) and as seen with other intrinsically disordered regions modelled by AlphaFold2 with the with the disordered region tending to be packed with the main domain (Ruff & Pappu, 2021). Indeed, the quality metrics for the AlphaFold2 Oca2 structure with the TMAlphaFold database highlight residues that are present in the membrane that should not be.

An AF2 remodeling exercise was performed in order to attempt to obtain a model where the N-terminal disordered region does not transverse the membrane boundaries as defined by the OPM server (Lomize, Pogozheva, Joo, Mosberg, & Lomize, 2012). The output provided 5 new models. The highest-ranking model with a pLDDT score of 72.9 showed the disordered loop mostly on the cytosolic side of the membrane with the exception of a 5-residue region (60-65) dipping into the membrane bi-layer (Figure 6).

**Figure 6.**
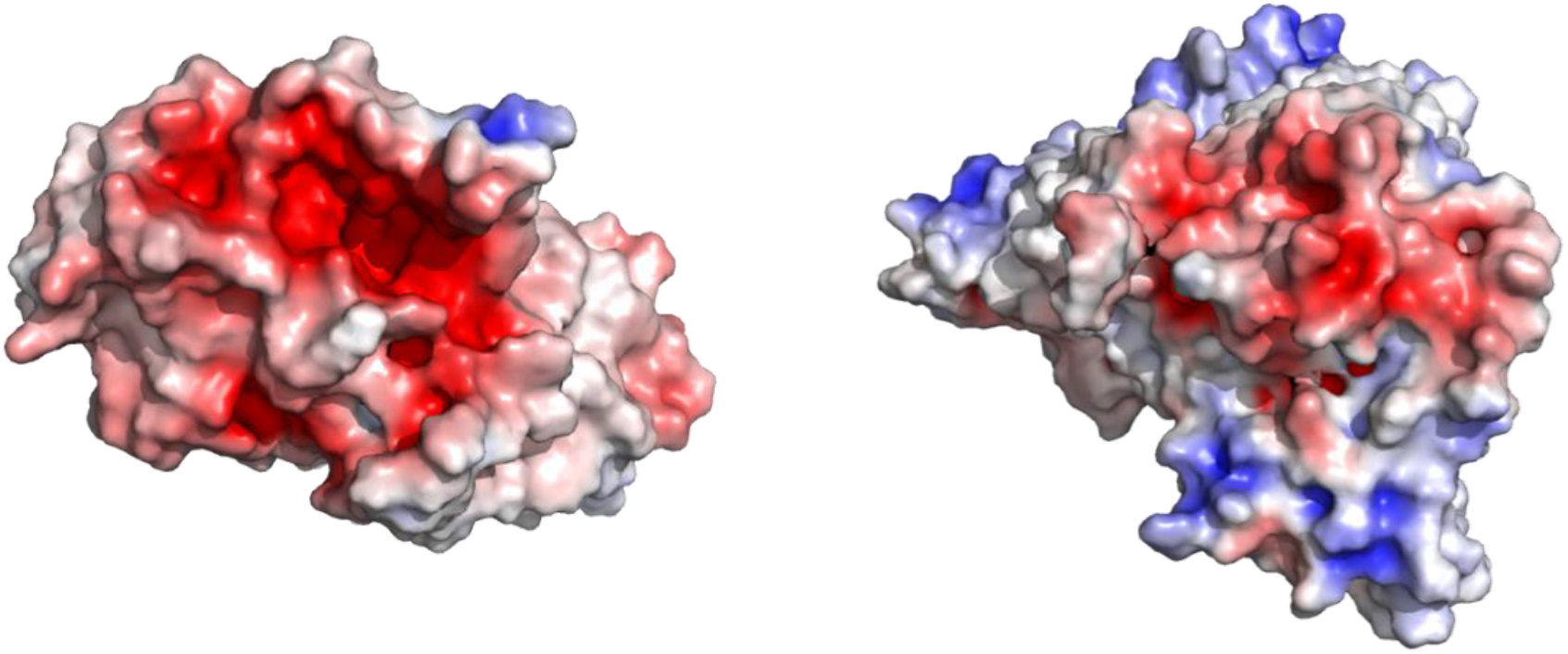
a) APBS electrostatic mapping surface view of Oca2 (transport domain only). b) APBS electrostatic mapping surface view of NaCT. Spectrum: Red (negative) to blue (positive).

Previous experimental studies have identified this loop to be cytoplasmic and possesses three dileucine motifs that required for human Oca2 function. These motifs have been shown to be essential for the targeting and localisation of Oca2 to the melanosome membranes by interacting with members of the clathrin-associated heterotetrameric adaptor protein family, AP-1 and/or AP-3 (Sitaram et al., 2009).

### Oca2 possesses a GOLD-like domain

The AlphaFold2 model predicts that the first luminal 130 residue loop forms an eight stranded beta sandwich. Examination of experimental PDB structures of other members of the IT superfamily revealed that Oca2 is the only member of the superfamily to possess this beta sandwich structure. Screening the 130-residue sequence of the beta sandwich against the PDB using HHpred did not yield any significant hits. In order to identify proteins possessing structurally similar regions, the 130-residue beta sandwich region was extracted from the AlphaFold2 model of Oca2 and screened, using Dali, against the full PDB. The results (Supplementary Table 2) gave a top hit with a Z-score of 10.3 for the central domain of tripeptidyl-peptidase 2 (TPP2) which is involved in the oligomerisation of TPP2 (Schönegge et al., 2012). The other top hits from the screen are for proteins possessing the Golgi Dynamics (GOLD) domain and have Z-scores above 9. Although this region shows no sequence similarity to known GOLD domains, the identification of homology through structural information, independent of sequence similarity, agrees with previous studies that indicate that GOLD domains have low sequence identity even between family members (Nagae et al., 2016).

The functions of the GOLD domain are largely unknown although there are indications that it is involved in the trafficking of proteins from the endoplasmic reticulum to other subcellular compartments (Nagae et al., 2016). Oca2 is known to become terminally glycosylated when it transits to a post-ER compartment to the Golgi (Sitaram et al., 2009). Human Oca2 has three evolutionary conserved consensus N-glycosylation sites (Asn 214, 218, and 273) (Sitaram et al., 2009) within the putative GOLD domain (Sitaram et al., 2009) and it has been demonstrated that some ER-resident proteins undergo GOLD domain N-glycosylation which is important for their trafficking between the ER and Golgi (Pastor-Cantizano et al., 2017) All of this suggests that the region previously termed the N-terminal luminal loop in fact encodes a GOLD domain involved in the trafficking of Oca2 from the ER to the Golgi prior to localisation at melanosomal membranes.

### Mutations in the Oca2 putative pore region results in severe albinism

The pseudo inverse repeat - made up of a broken transmembrane helix, an amphipathic helix and a re-entrant loop packing against the broken transmembrane helix - forms a substrate-binding chamber possessing two flanking re-entrant loops with the N-terminal half of the re-entrant loop containing highly conserved residues. Consurf (Ashkenazy et al., 2016) analysis highlights this region as highly conserved. Mutations in Oca2 disrupt melanin production within the melanosome. The hindered melanin synthesis has been linked to altered melanosome luminal pH which is correlated with reduced chloride conductance across the melanosome membrane (Bellono et al., 2014).

Positions of mutations that are characterised in vitro or in vivo were mapped onto the AlphaFold2 model in order to provide a structural context. Mutations at positions V443 and K614 are known to result in oculocutaneous albinism type II as well as inhibiting *in vitro* melanosome melanin content. Mapping these mutations onto the Alphafold2 model reveals that V443 is present on the N-terminal re-entrant loop of the first repeat unit and has a critical impact on chloride conductance across the melanosome membrane. The position of this functionally critical residue on a re-entrant loop is in accordance with other transporters where the re-entrant loop has a role in channel specificity (Mesdaghi, Murphy, Sánchez Rodríguez, Burgos-Mármol, & Rigden, 2021). K614 is present on the predicted cytoplasmic loop between the two inverse repeat units that would not contribute to any transport functionality of the protein; this is in agreement with the fact that in vitro studies show that K614 mutations have little effect on chloride conductance. K614 mutations present in albino patients also possess additional Oca2 mutations (Passmore, Kaesmann-Kellner, & Weber, 1999), so that the K614 change may itself not be of critical phenotypic importance.

Furthermore, mapping the 5-point mutation from Bellono et al (5mut: V443I, M446V, I473S, N476D, N489D) shows that they are all present on the N-terminal re-entrant loop/transmembrane helix structural motif of the first repeat. Again, these mutations result in inhibition of chloride conductance and are present in the putative pore region of the protein.

Mapping of other known mutations that result in severe albinism (F385, M395, N489, N679) (King et al., 2003; Passmore et al., 1999; Simeonov et al., 2013) show that these critical residues are present in either the N- or C-terminal transport domain region; F385 and M395 in the first transmembrane helix of the N-terminal transport domain region, N489 in the second transmembrane helix of the N-terminal transport domain region that packs with the re-entrant loop and W679 in the first transmembrane helix of the C-terminal transport domain region (Figure 5).

The mapping of known Oca2 mutations show that those present in the inverted repeat that forms the putative pore are more likely to result in the severe oculocutanous albinism type 2 phenotype. Similarly, mutation of Oca2 in regions important for melanosome localisation also results in the severe phenotype of oculocutanous albinism type 2. However, in contrast, mutations localised relatively distant to the putative pore region do not result in the severe phenotypes of oculocutanous albinism type 2.

### Citrate docks at the putative binding site

Bellono et al speculate that Oca2 might be an accessory subunit of a Cl^-^ transporter or form a Cl- channel or carrier protein itself like the bacterial homologue ArsB. The pseudo inverted repeat topology that includes re-entrant loops facing each other in the membrane packed against transmembrane helices has been seen previously in other chloride transporters such as CLC transporters (Feng, Campbell, Hsiung, & MacKinnon, 2010; Mesdaghi et al., 2021). Attempts were made to perform Oca2 docking a chloride ion. Docking of chloride was not successful; chloride did not dock at the putative binding site and was placed outside of the transport domain. Consequently, as Oca2 has strong HHpred hits with SLC13 transporters, docking of the SLC13 substrate was considered.

SLC13 transporters are members of the larger divalent-anion sodium symporter (DASS) family (Bergeron, Clémençon, Hediger, & Markovich, 2013; Markovich & Murer, 2004; Pajor, 2014; Prakash, Cooper, Singhi, & Saier, 2003). Most DASS transporters are sodium-coupled symporters that transport one substrate for each 2–4 sodium ions. However, some DASS members are antiporters (Pos, Dimroth, & Bott, 1998). Sequence analysis and examination of experimental models show that DASS symporters and antiporters share the same fold (Lolkema & Slotboom, 1998). The DASS antiporters possess surrogate residues (K,R or H) to compensate for the absence of sodium ions. Examination of Oca2 at the surrogate residue equivalent positions shows that the surrogate residues (K,R or H) that are present in the experimental structure of the DASS antiporter 6wu1 (LaINDY) are not present in Oca2. Furthermore, the substrate binding residues where the side chain is involved in the binding of two sodium ions identified in the DASS symporter experimental structure of 7jsk (NaCT) (Asn141 and Asn465) are present at the equivalent positions in the Oca2 model (Asn442 and Asn741). Indeed, visualization of the electrostatic surface view of Oca2 highlights a negatively charged region corresponding to the putative Na^+^ binding site (Figure 6).

In the human NaCT experimental structure (Sauer et al., 2021) a density which appears to be citrate is observed. There, in addition to Asn141 and Asn465, the authors propose additional substrate binding residues that form a substrate binding motif for citrate (Ser140-Asn141-Thr142 and Ser464-Asn465-Val466). The Oca2 model only possesses the Asn equivalents. Webina (Kochnev et al., 2020) was subsequently employed in an attempt to dock citrate at the equivalent position in Oca2 as seen in NaCT. The citrate does indeed dock on Oca2 at the equivalent position as observed in NaCT (Figure 7). Furthermore, docking of citrate to the AlphaFold2 model of NaCT results in docking again at the equivalent position with similar scores (ΔG −6.3 kcal/mol and −5.1 kcal/mol for NaCT and Oca2 respectively). To confirm this a rescoring exercise was performed for both Oca2 and NaCT; again, both NaCT and Oca2 obtained similar scores of ΔG −5.4 kcal/mol and −5.5 kcal/mol respectively. Attempting to dock citrate on to ArsB, however, does not result in docking of citrate at the putative binding site. Indeed, conservation mapping using Consurf highlights the putative ligand binding residues as highly conserved.

**Figure 7:**
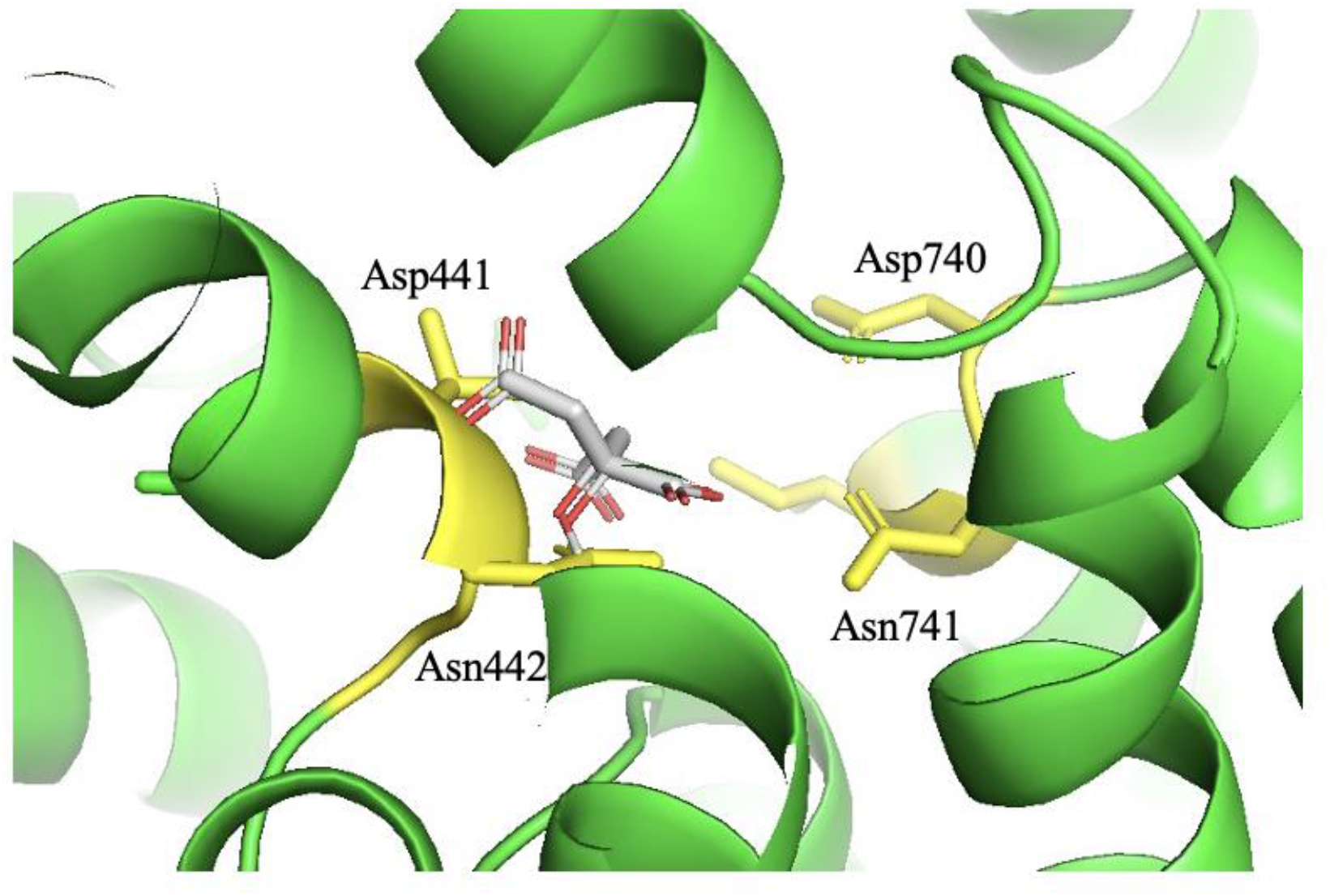
Webina Oca2 docking of citrate. Yellow are the conserved Asn442/Asn741 and adjacent residues.

These results suggest that citrate is plausible substrate for Oca2, but this does not align with the experimental observation that Oca2 is involved with chloride conductance across the melanosome membrane. Dicarboxylates are known to have a role in metabolic signaling. It is plausible that the movement of citrate (or another dicarboxylate) in and out of the melanosome could modulate chloride conductance across the melanosomal membrane downstream. Indeed, it has been shown that citrate inhibits melanin synthesis via the GSK3*β*/*β*-catenin signaling pathway which involves the regulation of tyrosinase transcription factors (Zhou & Sakamoto, 2020); citrate may be involved in the regulation of melanin synthesis at other key points in synthesis pathway.

### AlphaFold2 multimeric modelling protocol in combination with traditional homology modelling was able to model Oca2 in alternative conformations

Given that DASS proteins operate via an elevator-type transport mechanism (Drew & Boudker, 2016; Reyes, Ginter, & Boudker, 2009) and the obvious homology that Oca2 shares with DASS transporters it can be confidently predicted that the Oca2 transport mechanism is also of the elevator type. The elevator-type transport mechanism involves the sliding of the transport domain through the bilayer as a rigid body while the scaffold domain remains fixed in order to achieve the transitions between the outward- and inward-facing states (Garaeva & Slotboom, 2020). During the transport cycle it has been demonstrated that DASS symporters cotransport by binding sodium first and then their substrate, with the reverse occurring during the release (Hall & Pajor, 2005; Mulligan, Fitzgerald, Wang, & Mindell, 2014; Pajor, Sun, & Leung, 2013; Wright, Hirayama, Kaunitz, Kippen, & Wright, 1983; Yao & Pajor, 2000). During the course of transition between the outward and inward facing states the angle between the amphipathic helices and the re-entrant loops change by approximately 30° allowing the movement of the transport domain (Figure 8); the presence of flexible hinge loops between the amphipathic helices and the re-entrant loops facilitate transport domain movement (Sauer et al., 2021).

**Figure 8.**
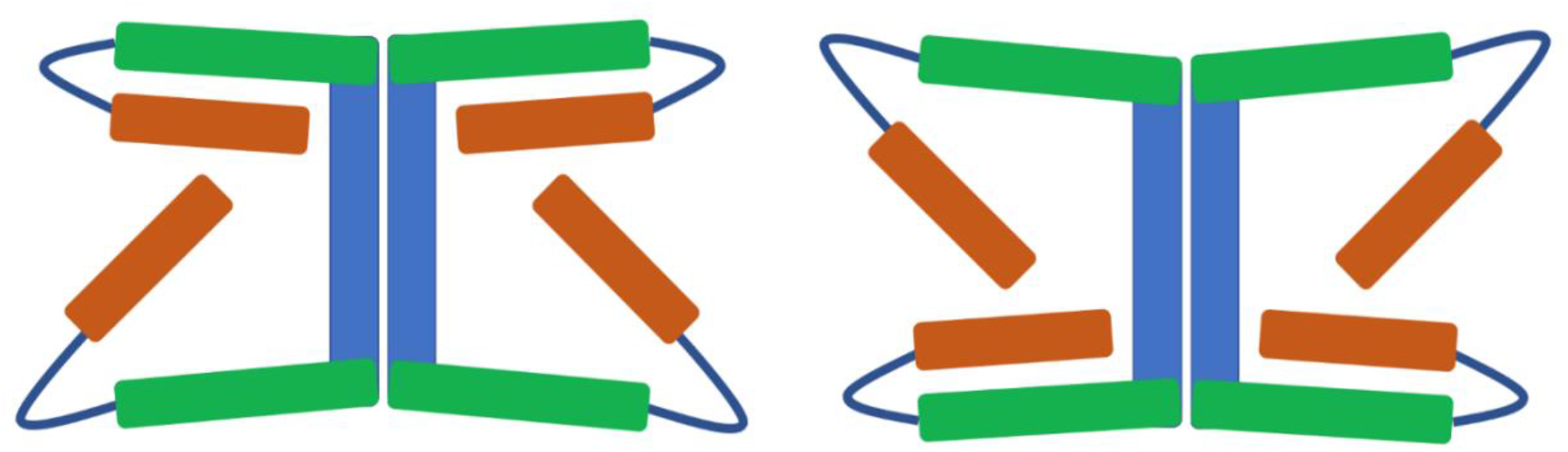
Blue: scaffold domain; Green: amphipathic helix; Orange: N-terminal of re-entrant loop. Between the inward and outward states, the angles at amphipathic (green)/re-entrant loop (only N-terminal side showing - orange) hinges (dark blue) change by around 30° resulting in the movement of the transport domain (not shown) relative to the scaffold domain.

Examination of the AlphaFold2 model showed that it was in the inward facing state where the angle between the N-terminal half of the re-entrant loop and the amphipathic helix is approximately 30° for the N-terminal side and approximately 55° for the C-terminal side. As observed for the DASS transporters VcINDY and LaINDY; the N-terminal angle increases by around 30° and the C-terminal angle decreases by around 30° when the transporter switches to the outward facing state (Sauer et al., 2021).

Attempts were made to model Oca2 in an outward facing conformation; building Oca2 locally using ColabFold resulted in 5 models all in the inward-facing conformation. Application of strategies that have previously been successful in the sampling of the conformational space of transporters were also employed; feeding ColabFold with shallow multiple sequence alignments (del Alamo, Sala, Mchaourab, & Meiler, 2022). Implementing this strategy also failed to generate models outside of the inward-facing conformational state. Further attempts were made to model Oca2 in an outward facing conformation by a combination of utilizing the VcINDY outward-facing structure as a template for AlphaFold2 (Mirdita et al., 2022) and by providing AlphaFold2 with reduced depth MSAs. However, again, AlphaFold2 was only able to generate the inward-facing conformation. The inability to obtain the outward facing conformation may be due to there being only one outward facing entry in the PDB with all others being a single DASS symporter in a substrate-bound, inward-facing state; resulting in modeling to converge on the inward facing state. Indeed, the AlphaFold2 prediction neural networks were trained on all structures deposited in the PDB on or before April 30, 2018 (Jumper et al., 2021) and many DASS homologues in the inward facing state were deposited before this date with the few outward facing examples being deposited in 2021 (Sauer et al., 2021).

The failure to generate Oca2 in an alternative by using previously known methods resulted in the employment of a novel strategy. A homology model of Oca2 was built using the VcINDY outward facing structure (6wtw) as a template for Modeller (Eswar et al., 2008). The output structure had a Dali alignment Z score with 6wtw of 54.8, however, the Oca2 structural features that are not present in VcINDY such as the putative GOLD domain were obviously not modelled. In order to rectify this, the Modeller model was then used in ColabFold in conjunction with various custom MSAs of varying depths to build a series of Oca2 structures (Table 4).

**Table 4.**
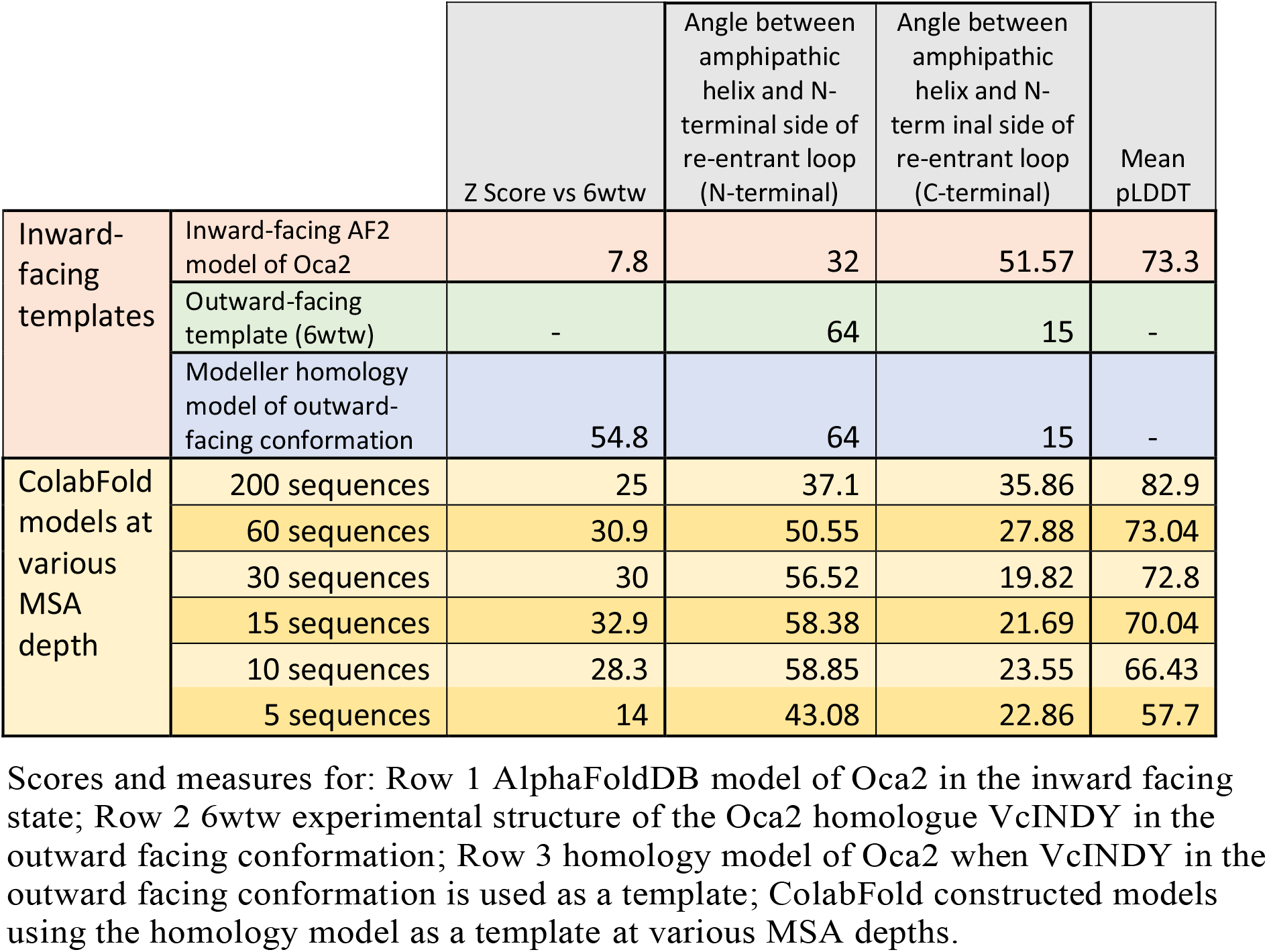
Z-scores, angles and quality scores for the Oca2 models.

|  |  | Z Score vs 6wtw | Angle between amphipathic helix and N-terminal side of re-entrant loop (N-terminal) | Angle between amphipathic helix and N-terminal side of re-entrant loop (C-terminal) | Mean pLDDT |
| --- | --- | --- | --- | --- | --- |
| Inward-facing templates | Inward-facing AF2 model of Oca2 | 7.8 | 32 | 51.57 | 73.3 |
|  | Outward-facing template (6wtw) | - | 64 | 15 | - |
|  | Modeller homology model of outward-facing conformation | 54.8 | 64 | 15 | - |
| ColabFold models at various MSA depth | 200 sequences | 25 | 37.1 | 35.86 | 82.9 |
|  | 60 sequences | 30.9 | 50.55 | 27.88 | 73.04 |
|  | 30 sequences | 30 | 56.52 | 19.82 | 72.8 |
|  | 15 sequences | 32.9 | 58.38 | 21.69 | 70.04 |
|  | 10 sequences | 28.3 | 58.85 | 23.55 | 66.43 |
|  | 5 sequences | 14 | 43.08 | 22.86 | 57.7 |

Scores and measures for: Row 1 AlphaFoldDB model of Oca2 in the inward facing state; Row 2 6wtw experimental structure of the Oca2 homologue VcINDY in the outward facing conformation; Row 3 homology model of Oca2 when VcINDY in the outward facing conformation is used as a template; ColabFold constructed models using the homology model as a template at various MSA depths.

The output models displayed the characteristic inverse repeat architectures of the scaffold and transport domains as seen in DASS transporters as well as the putative GOLD domain as predicted in the earlier modelling of the inward facing state of Oca2. Reducing the MSA depth did negatively influence the quality scores of the output models but at the same time improved the Dali alignment Z score with respect to the outward facing DASS transporter 6wtw thereby indicating more outward-facing structures. Additionally, the measurement of the N-terminal amphipathic/re-entrant hinge angle and C-terminal amphipathic/re-entrant hinge angle showed the characteristic angle combinations of the outward facing state; N-terminal angle being larger than the C-terminal angle (Figure 9).

**Figure 9.**
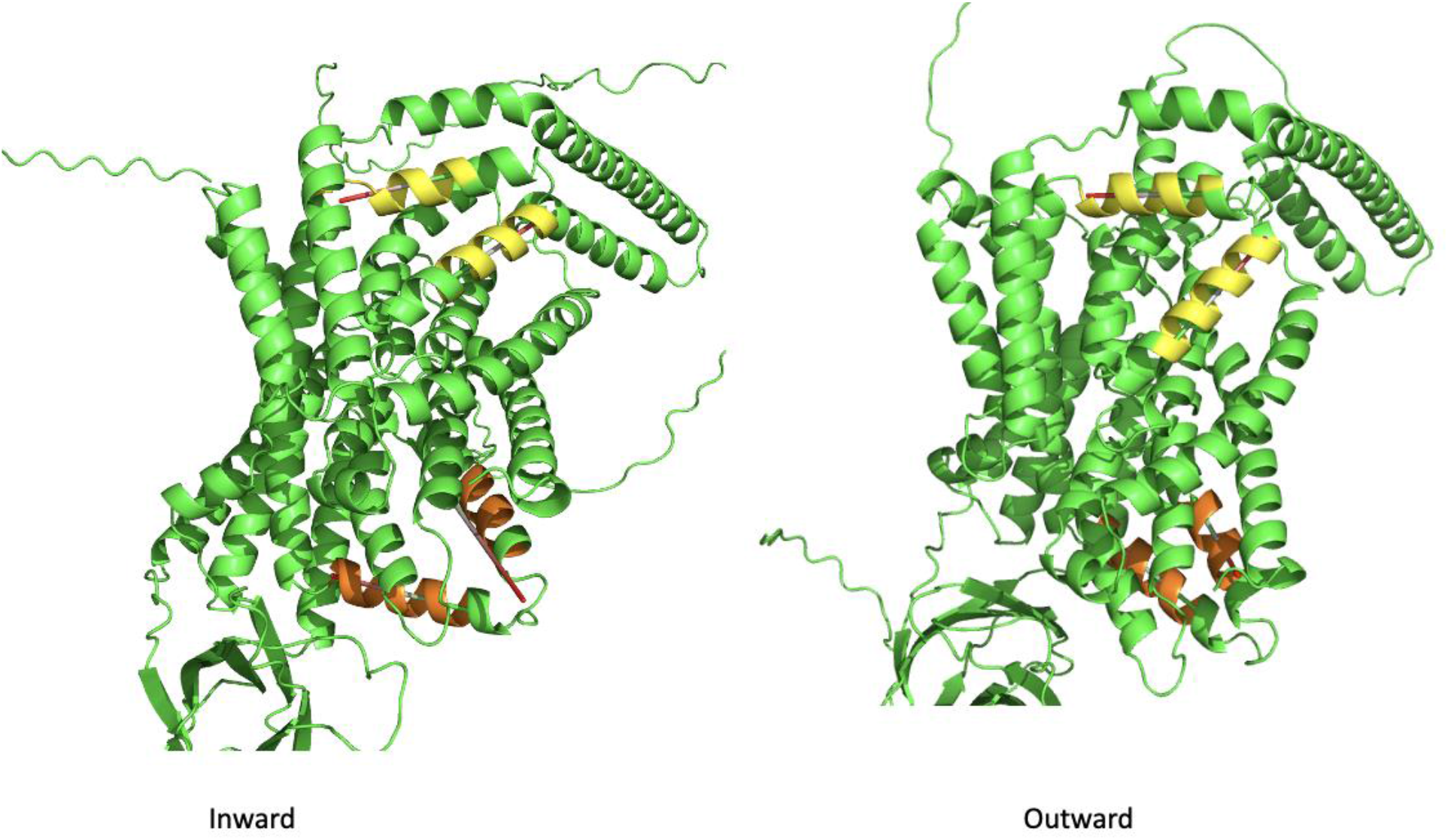
Yellow: N-terminal amphipathic/re-entrant hinge; Orange: C-terminal amphipathic/re-entrant hinge.

### AlphaFold2 is able to construct a plausible Oca2 homodimer in both conformations when used in combination with traditional homology modelling

The close homologues of Oca2 have experimental structures that form homodimers (Nie, Stark, Symersky, Kaplan, & Lu, 2017; Sauer et al., 2021, n.d.)/. Therefore attempts were made to model Oca2 as a homodimer. First, ColabFold was employed to build a homodimer without using a template and utilizing the full MSA. This produced five models each of which were in the inward facing conformation (Figure 11). Next the Modeller template and reduced MSA (15 sequences) were used to build a homodimer; this produced five models in the outward facing conformation (Figure 10).

**Figure 10.**
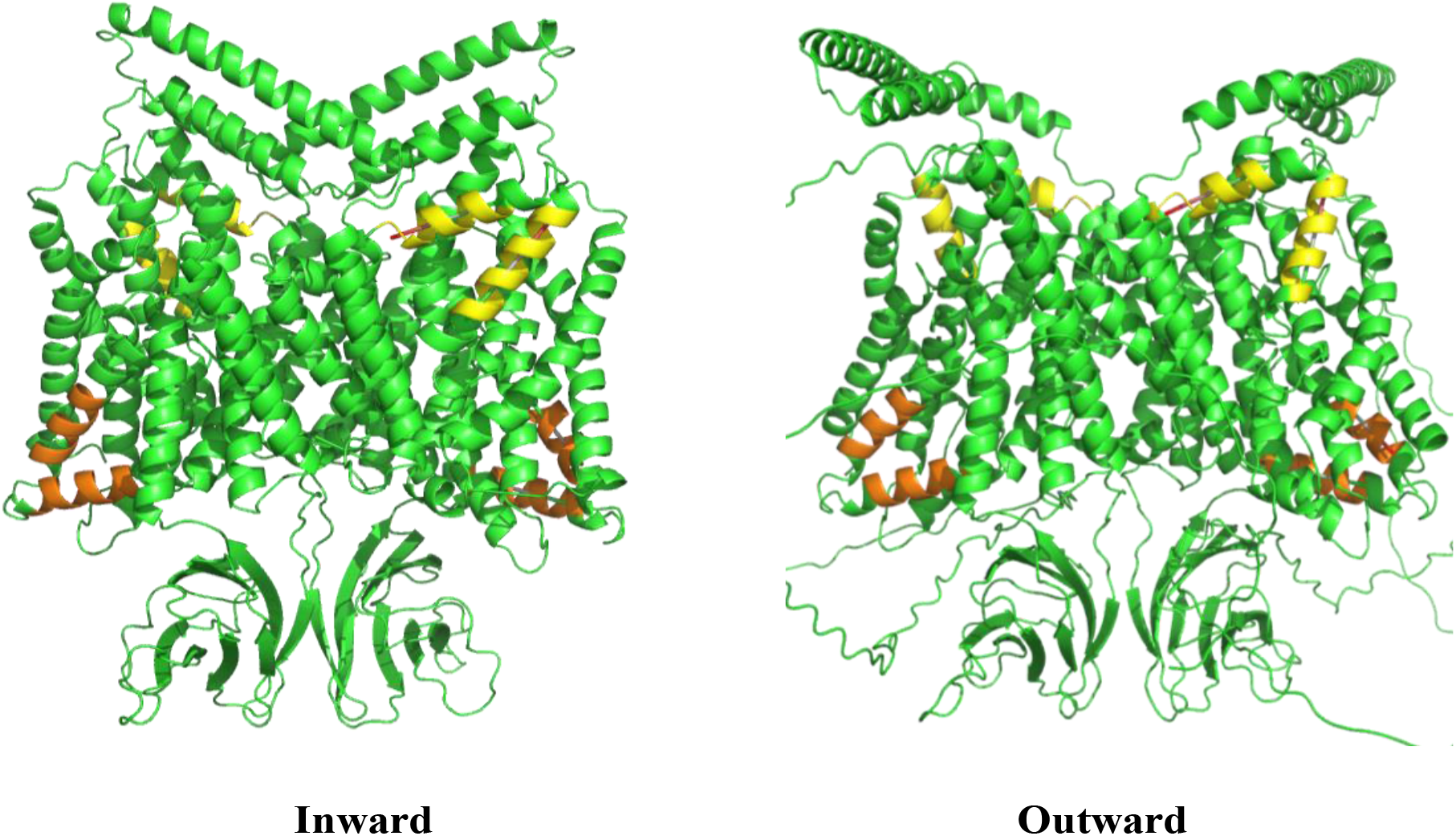
Yellow: N-terminal amphipathic/re-entrant hinge; Orange: C-terminal amphipathic/re-entrant hinge.

**Figure 11:**
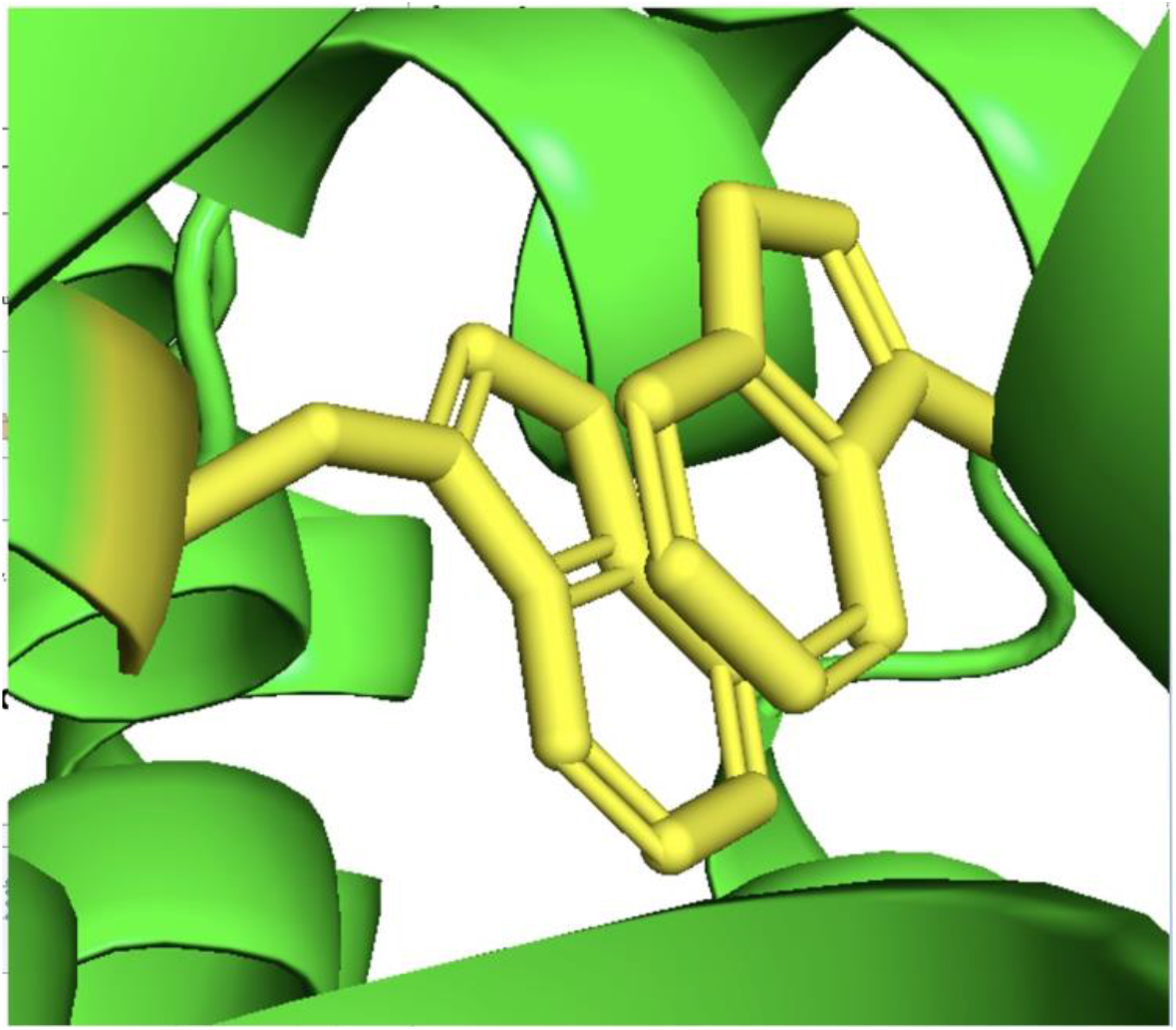
Pi-Pi interaction

Pisa (Krissinel & Henrick, 2007) was employed to determine the area of the interface which was calculated as 55111.2Å^2^ for both models. This interface area is much larger than NaCT; 21035.3 Å^2^. The presence of the putative GOLD domains in Oca2 contribute to this dimerization surface area resulting in this unusually large interface. To our knowledge, dimerized GOLD domains have not been reported previously. Further analysis of the homodimer interfaces of Oca2 revealed the presence of an interaction between Trp679 of both monomers which could contribute to the stabilisation of the interface formed from the two scaffold domains resulting in a stable rigid structure in the membrane (Figure 11); this same interaction can be seen in the dimer interface of NaCT; Pi–Pi interaction between Trp408 and Trp408’ from the neighbouring protomers stabilizing the two scaffold domains together into a rigid framework (Sauer et al., 2021).

## Conclusions

Oca2 shows structural similarities to SLC13 proteins. The AlphaFold2 model has provided the opportunity to revise the current consensus view of its topology. The AlphaFold2 model of Oca2 strongly suggests that it shares the DASS family topology and possesses a GOLD-like domain. The DASS family contains both symporters and antiporters; Oca2 possesses symporter features. Although the molecular specificity of Oca2 remains unclear, Oca2 possesses key citrate-binding residues as seen in NaCT and citrate docks to Oca2 at the putative binding site as observed in NaCT. When clinically relevant and in vitro mutations are mapped on to the model it is seen that they cluster on the transport domain of the structure. AlphaFold2 modelling of Oca2 has demonstrated that, like DAS transporters, Oca2 can exist in two plausible conformations: inward-and outward-facing, supporting an elevator-type transport mechanism.

## Supplementary

**Supplementary Table 1:**
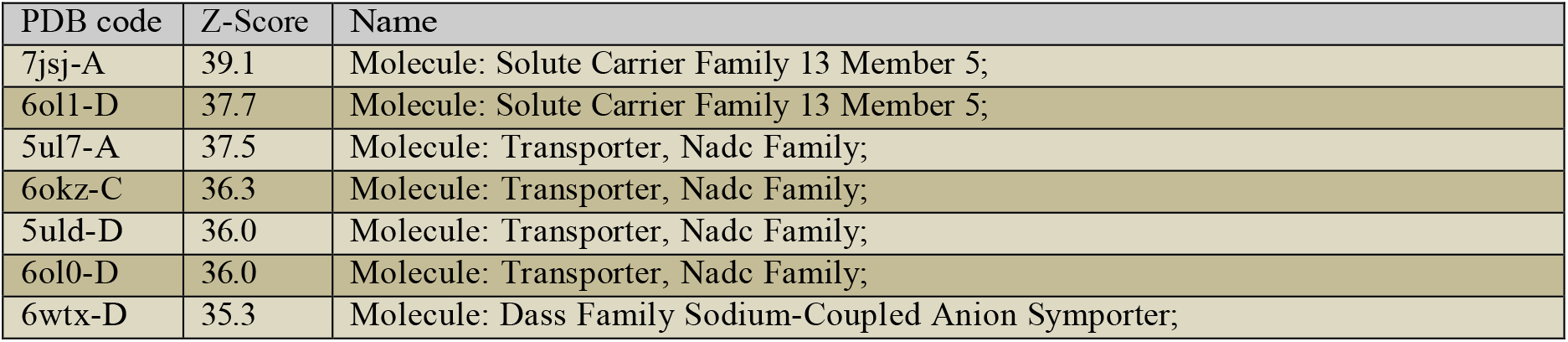
Dali results for structural screen of Oca2 against PDB

**Supplementary Table 2:**
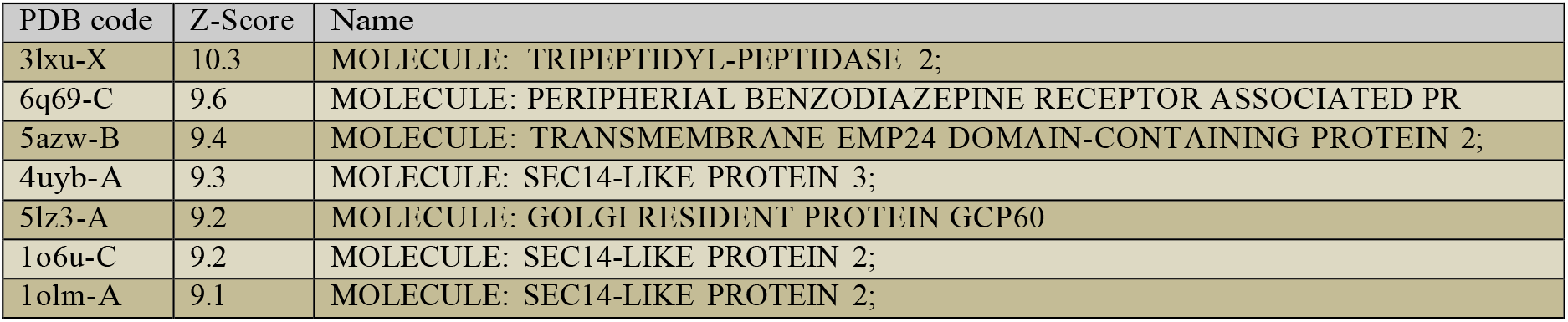
Dali results for structural screen of beta sandwich (residues 196-331) region against PDB

## Notes

### Competing Interest Statement

The authors have declared no competing interest.

