## Supplemental Table 1 and 2 for "Structural Insights into Pink-eyed Dilution Protein (Oca2)"

### Supplementary

Supplementary Table 1: Dali results for structural screen of Oca2 against PDB

| PDB code | Z-Score | Name |
|---|---|---|
| 7jsj-A | 39.1 | Molecule: Solute Carrier Family 13 Member 5; |
| 6ol1-D | 37.7 | Molecule: Solute Carrier Family 13 Member 5; |
| 5ul7-A | 37.5 | Molecule: Transporter, Nadc Family; |
| 6okz-C | 36.3 | Molecule: Transporter, Nadc Family; |
| 5uld-D | 36.0 | Molecule: Transporter, Nadc Family; |
| 6ol0-D | 36.0 | Molecule: Transporter, Nadc Family; |
| 6wtx-D | 35.3 | Molecule: Dass Family Sodium-Coupled Anion Symporter; |

Supplementary Table 2: Dali results for structural screen of beta sandwich (residues 196-331) region against PDB

| PDB code | Z-Score | Name |
|---|---|---|
| 3lxu-X | 10.3 | MOLECULE: TRIPEPTIDYL-PEPTIDASE 2; |
| 6q69-C | 9.6 | MOLECULE: PERIPHERIAL BENZODIAZEPINE RECEPTOR ASSOCIATED PR |
| 5azw-B | 9.4 | MOLECULE: TRANSMEMBRANE EMP24 DOMAIN-CONTAINING PROTEIN 2; |
| 4uyb-A | 9.3 | MOLECULE: SEC14-LIKE PROTEIN 3; |
| 5lz3-A | 9.2 | MOLECULE: GOLGI RESIDENT PROTEIN GCP60 |
| 1o6u-C | 9.2 | MOLECULE: SEC14-LIKE PROTEIN 2; |
| 1olm-A | 9.1 | MOLECULE: SEC14-LIKE PROTEIN 2; |
